# Increased activity of IRE1 improves the clinical presentation of EAE

**DOI:** 10.1101/2023.04.19.537391

**Authors:** Valerie Bracchi-Ricard, Kayla Nguyen, Daniela Ricci, Brian Gaudette, Jorge Henao-Meija, Roberta Brambilla, Tetyana Martynyuk, Tali Gidalevitz, David Allman, John R. Bethea, Yair Argon

## Abstract

Activation of the ER stress sensor IRE1α contributes to neuronal development and is known to induce neuronal remodeling *in vitro* and *in vivo*. On the other hand, excessive IRE1 activity is often detrimental and may contribute to neurodegeneration. To determine the consequences of increased activation of IRE1α, we used a mouse model expressing a C148S variant of IRE1α with increased and sustained activation. Surprisingly, the mutation did not affect the differentiation of highly secretory antibody-producing cells, but exhibited a strong protective effect in a mouse model of experimental autoimmune encephalomyelitis (EAE). Significant improvement in motor function was found in IRE1C148S mice with EAE relative to WT mice. Coincident with this improvement, there was reduced microgliosis in the spinal cord of IRE1C148S mice, with reduced expression of pro-inflammatory cytokine genes. This was accompanied by reduced axonal degeneration and enhanced CNPase levels, suggestiing improved myelin integrity. Interestingly, while the IRE1C148S mutation is expressed in all cells, the reduction in proinflammatory cytokines and in the activation of microglial activation marker IBA1, along with preservation of phagocytic gene expression, all point to microglia as the cell type contributing to the clinical improvement in IRE1C148S animals. Our data suggest that sustained increase in IRE1α activity can be protective *in vivo*, and that this protection is cell type and context dependent. Considering the overwhelming but conflicting evidence for the role of the ER stress in neurological diseases, a better understanding of the function of ER stress sensors in physiological contexts is clearly needed.

## Introduction

Inositol requiring enzyme 1 alpha (IRE1α) is the ancient and most conserved stress sensor that initiates the unfolded protein response (UPR). When activated, IRE1α cleaves the transcript XBP1 and removes a short intron (1). The two remaining RNA ends require non-conventional ligation (2), mediated by the RNA 2’,3’-cyclic phosphate and 5’-OH ligase RtcB, (3), creating spliced XBP1 (XBP1s). XBP1s encodes an active basic-leucine zipper transcription factor (1), which regulates the expression of hundreds of stress responsive genes, such as chaperones (e.g. Grp78/BiP) and lipid biosynthesis genes for ER expansion (4, 5). Inability of the UPR to re-establish protein homeostasis following stress is linked to several neurodegenerative diseases such as Alzheimer’s or Parkinson’s diseases (reviewed in (6)). The UPR is also an important regulator of neuroinflammation (7) and is upregulated in experimental autoimmune encephalomyelitis (EAE), a demyelinating disease that affects nerve conduction and leads to paralysis and serves as a common mouse model for multiple sclerosis (MS), (reviewed in (5)) which currently has no cure. The disease is mediated by T cells that autoreact to myelin components and cause inflammation in the central nervous system (8). Because several UPR proteins are upregulated in MS and EAE lesions (9, 10), it stands to reason that modulating the UPR response may be a useful therapeutic intervention. Yet, it is unclear whether upregulation of the UPR seen in neurodegeneration, including in MS or EAE, represents an adaptive response that is insufficient, or an overactivated response with a known potential to be maladaptive. The maladaptive potential of overactivated IRE1 pathway is linked to pro-apoptotic signaling (11). Despite that, selective activation of the IRE1α pathway can be protective in stress conditions (12). Indeed, ectopic overexpression of spliced XBP1 was neuroprotective in a model of EAE/optic neuritis when coupled with inhibition of PERK pathway (13). The goal of our study was to determine whether increasing activation of IRE1 in a physiological context has a protective effect in the mouse EAE model. We generated a mouse carrying a mutant form of IRE1α, namely IRE1C148S, that exhibits enhanced and prolonged activity (14), with the prediction that enhancing IRE1α activity would benefit cells with high secretory loads, such as oligodendrocytes and neurons in the CNS and antibody-secreting B cells in the periphery, by increasing their secretory capacities. To our surprise, plasma cells from IRE1C148S had the same antibody secretion capabilities as their WT counterparts. However, enhanced IRE1α activity was beneficial for reducing the symptoms of EAE in mice, showing potential for IRE1α activation in the treatment of MS.

### Key results

- Generation and characterization of a novel IRE1C148S variant mouse with activity-enhancing mutation.
- Improved clinical outcome in IRE1C148S mutant mice following EAE.
- Cell-type specific effect of hyperactive IRE1α, with plasma cells being unaffected.
- Reduction in microgliosis and preserved CNPase levels in IRE1C148S mutant mice correlate with lower clinical scores.
- Reduction in proinflammatory cytokine levels but preservation of phagocytosis gene expression.
- Increased MBP production but no difference in proliferation or differentiation of IRE1C148S mutant oligodendrocyte precursor cells

## Materials and Methods

### Mice

IRE1α C148S mice were generated using the CRISPR technology in the Transgenic and Chimeric Mouse facility of the Department of Genetics at University of Pennsylvania (Philadelphia, PA). C57Bl/6J mice were obtained from Jackson’s lab (stock# 000664). Mice were acclimated to Drexel University animal facility for at least 1 week before any experimental procedure. Mice were housed in groups of up to 5 mice in a humidity/temperature-controlled environment with 12h daylight cycle along with access to food and water *ad libitum*. All animal experiments were approved by Drexel University’s Institutional Animal Care and Use Committee.

### Genotyping the IRE1C148S mutation

DNA was extracted from tail biopsies using the alkaline lysis method. PCR was performed using primers *genC148S_seq_Fw: 5’- ACA TCC TGG CAT TTC AGG -3’* and *genC148S_seq_Rv: 5’- AGC TGT AGG TAG TGC ACC -3’.* The purified PCR product (Qiaquick PCR purification kit, Qiagen) was sequenced using the forward primer (Genewiz, South Plainfield, New Jersey).

### RNA extraction and quantitative real-time PCR

Total RNA from lumbar spinal cords from WT and IRE1C148S 31d post-EAE (and controls) were extracted using TRIzol™ (ThermoFisher Scientific) and further purified on RNAeasy mini columns using on column DNA digestion according to manufacturer’s protocol (Qiagen). For each sample, an aliquot of RNA (500 ng) was reverse transcribed using the Ominiscript RT kit (Qiagen). Real-time PCR quantification was performed using Sybr green and primers specific for the gene of interest, using the standard curve method, where a standard curve is run with known amounts of the target PCR product, and the experimental sample concentrations are determined based on the standard curve. Every RT reaction is also subjected to amplification with β-actin primers as a control, and all data are normalized to β-actin.

### qRT-PCR for XBP1s

Total RNA was isolated with the Trizol reagent (Thermo FisherScientific) following manufacturer’s instructions. Two hundred ng of RNA were retrotranscribed to cDNA by priming with oligo(dT)_12–18_ and Superscript II retrotranscriptase (ThermoFisher Scientific). Quantitative PCR was performed using SYBR green reagent (Thermo Fisher Scientific) and the reaction run on Applied Bio-systems StepOne Plus machine. Data were analyzed using ΔΔCt method.Quantitative PCR primers: Rpl19 (Ribosomal Protein L19): forward: 5’- AAAACAAGCGGATTCTCATGGA-3’, reverse: 5’-TGCGTGCTTCCTTGGTCTTAG-3’; XBP1s: forward: 5’-CTGAGTCCGCAGCAGGTGCAG-3’, reverse: 5’-ATCCATGGGAAGATGTTCTGG- 3’.

### Western blot analysis

Proteins were extracted from different brain regions (cortex, hippocampus, cerebellum) and spinal cord of WT and IRE1C148S mutant mice as previously described (15). Equal amounts of each sample (20 Dg) were loaded on an 8% or 15% SDS-PAGE depending on the size of the protein to be detected. After transfer to nitrocellulose membrane using the TurboBlot system (Biorad), primary antibodies were applied: rabbit anti-CNPase (1:1,000, Cell signaling), rat anti-MBP (1:40.000, Millipore), mouse anti-NF200 (1:1,000, Sigma), rabbit anti-IBA1 (1:500, Cell signaling). Species-specific HRP-conjugated secondary antibodies from ImmunoJax were used. Following incubation with West Pico ECL substrate (Thermo Fisher Scientific), blots were imaged using the Gel Doc XR+ (BioRad) and analyzed with image J software (NIH). Protein quantification was normalized to the total protein for each sample determined after transfer to the membrane using Ponceau S solution (Sigma).

### EAE induction

Induction of experimental autoimmune encephalomyelitis was achieved by subcutaneous immunization of adult mice with an emulsion of MOG_35-55_ (Bio-Synthesis Inc.) peptide in complete Freund’s adjuvant containing heat-inactivated Mycobacterium tuberculosis (MT) H37Ra as previously described by (16). Each mouse received 2 s.c. injections of 50μl each into the posterior right and left flanks (total of 100μg of MOG and 200 μg of MT per mouse). A second injection of MOG/CFA was administered 7days later. Mice were monitored daily for clinical symptoms and body weight loss. Clinical scores were as follows: 0-no motor deficits, 1- tip of the tail is flaccid, 2- Tail completely flaccid, 3- Hindlimb paralysis or ataxia, impaired balance/ambulation, 4- Forelimb paralysis, 5- Moribund.

### Pain testing

Mechanical allodynia measurement was performed using the Von Frey test as previously described (17). In brief, filaments of different sizes (0.02g-2g) are applied to the hind paw of the mouse until a withdrawal response is elicited. Only when the withdrawal response is paired with cognitive awareness is the response considered as painful.

### Oligodendrocyte cultures and *in vitro* studies

Oligodendrocyte precursor cells (OPC) were isolated from post-natal day 3 WT and IRE1C148S pups using PDGRα magnetic beads (Miltenyi) with the neural dissociation kit (Miltenyi). OPCs were plated onto PDL-laminin coated plates. After the first passage, OPCs were plated onto PDL-laminin coverslips in 24-well plate at (20,000 cells/well) in proliferation medium (DMEM/F12 containing 2% B27, 1% N2, 20 ng/ml FGF2, and 10 ng/ml PDGFaa and 1% Pen-Strep). Differentiation into mature oligodendrocyte was induced by switching the culture medium to a medium lacking growth factor and containing thyroid hormone (T3, 40 ng/ml). Five to 7-days later the cells were fixed with 4% PFA and immunostained for MBP and CC1. IRE1α inhibitors A106 (50μM, Millipore) and 4μ8c (30μM, Millipore) were added to the differentiation medium. DMSO was used as the vehicle control.

### Electron microscopy and G-ratio

WT and IRE1C148S naïve or EAE d31 mice were deeply anesthetized with ketamine and xylazine prior to perfusion with PBS followed by 4% PFA in PBS. Spinal cords were dissected out then transferred to 2% glutaraldehyde, 100 mM sucrose in 0.05M Sorensen’s phosphate buffer, and sent to the Transmission Electron Microscopy Core facility of the Miami Project to Cure Paralysis (University of Miami Miller School of Medicine) for processing. The G-ratio, a measure of myelin thickness and compaction, calculated as fiber (axon + surrounding myelin) diameter/axon diameter, was determined as previously described (18).

### B-cell isolation and *in vitro* differentiation

Spleens were disrupted between two frosted slides and filtered through a nitex mesh, followed by red blood cell lysis with ACK lysis buffer (19). Splenocytes from 10-week-old mice were plated at 1 x 10^6^ cells/ml in 96-well culture plates in complete growth media (RPMI, 10% FBS, HEPES, Pen/Strep, NEAA, Na-Pyruvate, 2-mercaptoethanol). Cultures were treated with LPS 0.1 mM and IL5 (100 ng/ml) where indicated and incubated at 37°C / 5% CO_2_ for the indicated times.

### Flow cytometry

Staining of lymphocytes and flow cytometry was performed as described in (20), using a cocktail of appropriate antibodies at optimized concentrations for 30 min on ice. Flow cytometry was performed using BD LSRII and LSR Fortessa analyzers and cell sorting with BD FACS Aria cell sorter. Data analysis was performed using Flowjo 8.8.7 software.

### Elispot

Antibody secretion was assayed with Elispot plates as described in (19)

### Statistical analysis

Statistical analyses were performed with GraphPad Prism 6 software. For the comparison of two groups the two-tailed T-test was used. For more than 2 groups, One-Way ANOVA followed by Fischer post-hoc test for multiple comparison was used. For the behavior, a Two-way ANOVA was performed. A result is considered significant when p<0.05.

## Results

### Generation and characterization of a novel IRE1C148S variant mouse with activity-enhancing mutation

The studies described here were designed to determine if augmenting the physiological ER stress response affects an autoimmune demyelinating disease. To this end, IRE1C148S mutant mouse was generated using CRISPR technology to introduce a point mutation in the IRE1α gene, replacing the highly conserved cysteine 148 (murine position - C150) with a serine residue (Fig. 1A). The cysteine 148 is involved in the activation of IRE1α and is also the target of protein disulfide isomerase family A member 6 (PDIA6), an oxidoreductase that attenuates IRE1α activity (14). Because of the mutation, IRE1α activity is prolonged *in vitro*, in 293T cells, leading to more robust splicing of XBP1, an immediate substrate of IRE1 (21). *In vivo* as well, the level of XBP1s is increased in the spinal cord of mutant mice, as shown by quantitative RT-PCR (Fig. 1B), with no change in the levels of IRE1α protein (Fig. 1C). Increased XBP1s was also detected in several brain regions (Sup. Fig. 1). Therefore, these data confirm that *in vivo* as well as in cell culture the Cys148-to-Ser mutation increases IRE1α activity.

**Fig. 1.**
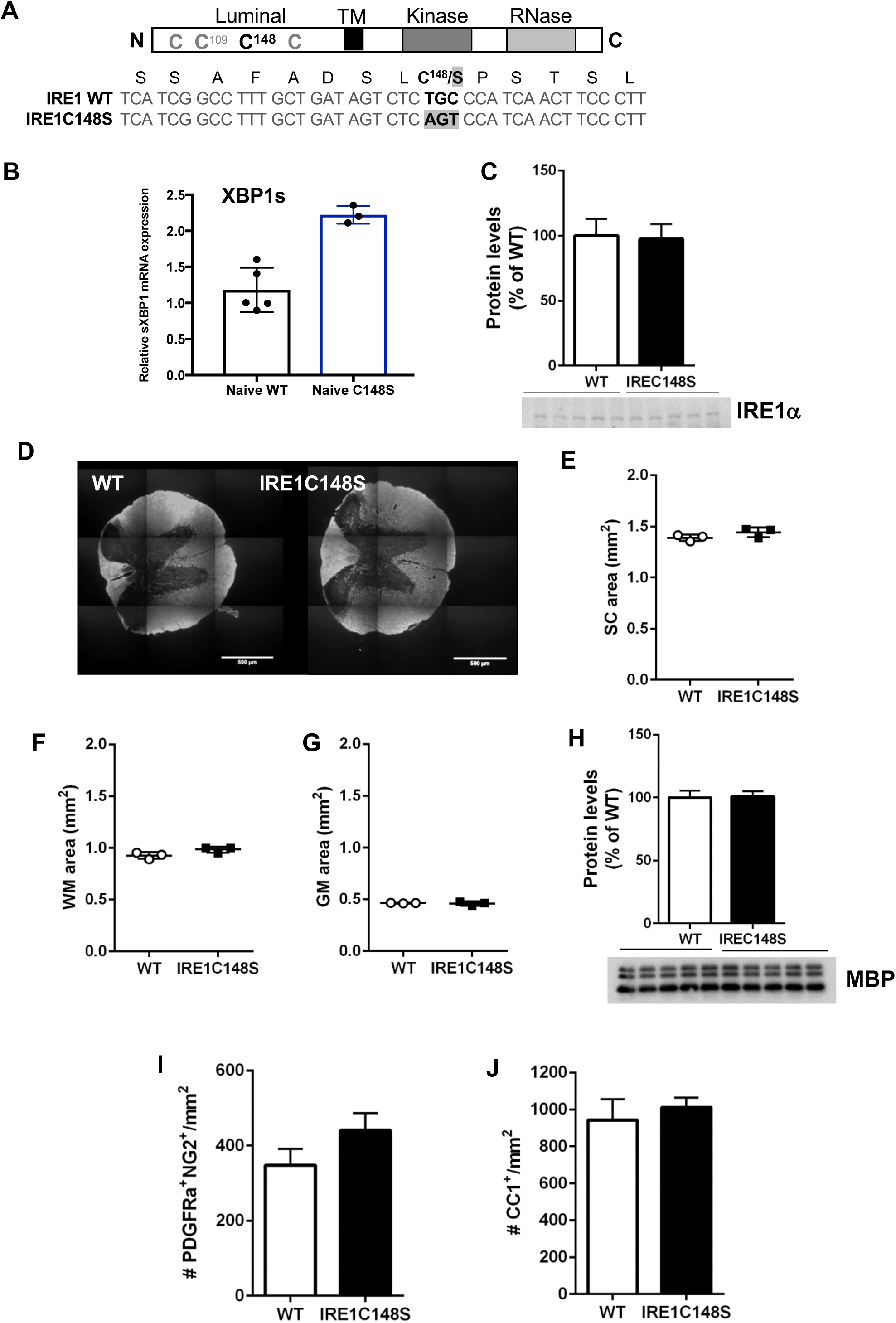
Spinal cord from naïve IRE1C148S variant with activity-enhancing mutation is not significantly different from WT mice. **A**- Schematic diagram of IRE1α protein and its main domains. The N-terminal domain resides in the lumen of the ER and serves as the stress sensing device. The four cysteine residues in this domain are indicated and the sequence encoding amino acids around Cys148 is listed below the diagram. TM, the transmembrane region that spans the ER membrane and is responsive to lipid stress (55). **B**- Quantitation of XBP1s transcripts in spinal cords of WT and IRE1C148S mice, using q-PCR. **C**- Quantification of IRE1α protein levels in the spinal cords of WT and IRE1C148S mice by Western blot. n=5/group. **D**- Representative thoracic spinal cord sections from WT and IRE1C148S mice immunostained for myelin basic protein (MBP). **E**- Spinal cord area quantification in WT and IRE1C148S mice. n=3/group **F-G**- Quantification of spinal cord areas: White matter (WM) area (**E**) and Grey matter (GM) area (**F**), n=3/group **H**- Western blot analysis of spinal cord extracts from WT and IRE1C148S for MBP. n=5/group **I**- Quantification of the number of oligodendrocyte precursor cells (PDGFRα^+^NG2^+^-cells) per mm^2^ in the spinal cord of WT and IRE1C148S. n=3/group **J**-Quantification of the number of oligodendrocytes per mm^2^ (CC1^+^-cells). n=3/group

Phenotypically, the mutant mice were indistinguishable from WT mice. Gross morphology of the spinal cord appeared normal, and we found no significant changes in white and grey matter between mutant and WT mice, either histologically or by myelin basic protein (MBP) expression (Fig. 1D-H). We further quantified the number of oligodendrocyte precursor cells (OPCs) (Fig. 1I) and oligodendrocytes (Fig. 1J) in the spinal cord, and found no significant differences.

### The IRE1C148S mutation improves clinical outcome in mice following EAE

Since no significant difference was observed between WT and IRE1C148S mice under normal physiological conditions, we sought to determine if a difference becomes evident under conditions that trigger IRE1α activation, such as in the course of the autoimmune demyelinating disease model EAE. A striking phenotype was found when EAE was induced by injection of MOG_35-55_ peptide in complete Freund’s adjuvant (but omitting the pertussis toxin as previously described (15)). As shown in Fig. 2A-C, the day of onset, defined as the first day when mice have a clinical score of 2 for two consecutive days (22), occurred significantly later for the IRE1C148S mutant mice than for control mice. Although differences between the groups in either the day of peak disease or in the severity score at peak disease did not reach statistical significance, suggesting that IRE1C148S mice do develop EAE similarly to control mice, the majority of IRE1C148S animals never exceeded the clinical score of 2, while the WT animals stayed at scores above 2 from day 19 until the end of experiment on day 31. Indeed, the extent of the disease was significantly reduced in IRE1C148S, as shown by the area under the curve (Fig. 2B), and the cumulative disease index (35 vs. 25, p<0.01, Fig. 2C). Furthermore, unlike WT mice, IRE1C148S mutants showed a significant improvement in the clinical score by the end of the experiment (day 31) in comparison to the peak disease (Fig. 2A). The IRE1C148S mice tended to lose less weight than control mice during the acute phase of the disease between days 18 and 20 (Fig. 2D). Thus, by several clinical criteria, the mutant IRE1C148S had a strong protective effect against EAE. However, mechanical allodynia assessment using Von Frey testing showed that both WT and IRE1C148S mutant mice developed pain which persisted for the duration of the study, with no statistical significance between groups (Fig. 2E).

**Fig. 2.**
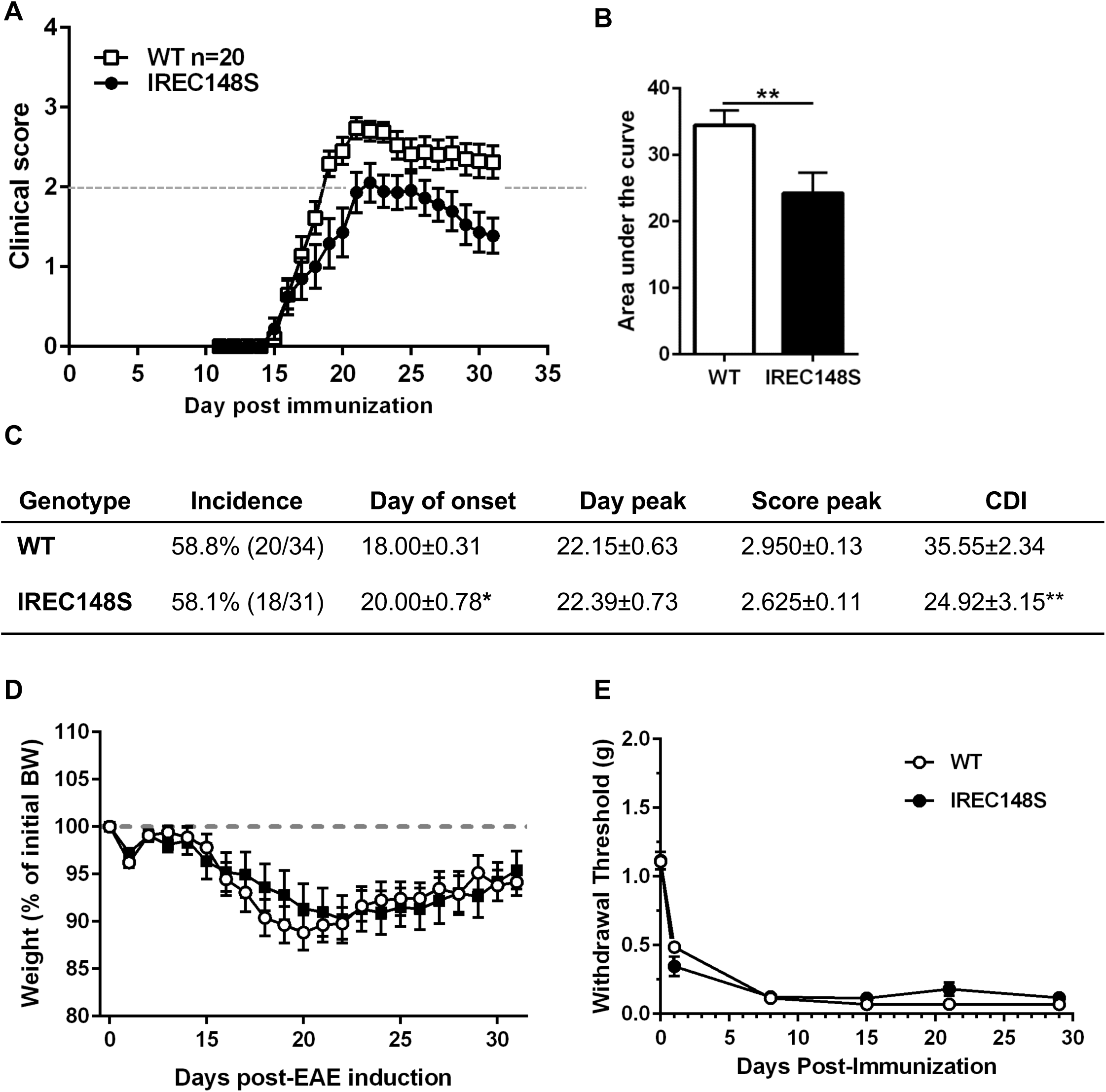
IRE1C148S rescues clinical phenotypes in the EAE model. **A**- Mice were MOG-immunized on day 0 and day 7 and monitored daily for motor clinical score until day 31 post immunization. n=20 WT, n=18 IRE1C148S. **B**- IRE1C148S mutant mice show a significant reduction in disease severity as measured by the area under the curve (AUC) compared to WT mice. n=20 WT, n=18 IRE1C148S. **C**- Table comparing different parameters of EAE between WT and IRE1α mutant mice. The day of onset is defined as the first day when a mouse has a clinical score of 2 for two consecutive days. The cumulative disease index (CDI) was calculated as the sum of the clinical scores between d7 and d31. **D**- Body weight (BW) was recorded daily after the second immunization and plotted as a percentage of initial BW. The graph shows the average ± sem of n=20 WT and 18 IRE1C148S with motor deficits. **E**- Mechanical allodynia in WT and IRE1C148S mice following EAE induction were determined using the Von Frey method. n=15 WT and n=10 IRE1C148S

### Enhanced IRE1 activity reduces myelin pathology and neuronal damage

To begin investigating changes that would help explain improvements in functional recovery, we analyzed myelin pathology and neuronal damage. To this end, we utilized EM to analyze naïve and diseased spinal cord tissue.

G-ratio analysis was used to investigate myelin thickness of axons between 0.5 and 2μm in diameter in diseased WT and C148Smice relative to naïve controls (Fig. 3A). There was no difference in the G-ratios between naïve and diseased IRE1C148S mice, indicating little to no detectable myelin degeneration in the mutants (Fig.3B). However, there was a significant reduction in G-ratios of diseased WT relative to naïve controls Similar results could also be observed by staining spinal cord lumbar sections from WT and IRE1C148S with the myelin marker MBP, with noticeable areas of demyelination in WT but not mutant mice (Fig. 3C). Further, we observed reduced axonal degeneration in diseased IRE1C148S mice relative to diseased WT mice (Fig. 3D). Western blot analysis demonstrated a significant increase in 2’, 3’-cyclic nucleotide 3’-phosphodiesterase (CNPase), an abundant protein in oligodendrocytes (Fig. 3E). Similarly, there was increased level of neurofilament 200 (Fig. 3F), which plays a vital role in organizing neuronal cytoarchitecture, in diseased IRE1C148S mice relative to diseased WT mice.

**Fig. 3.**
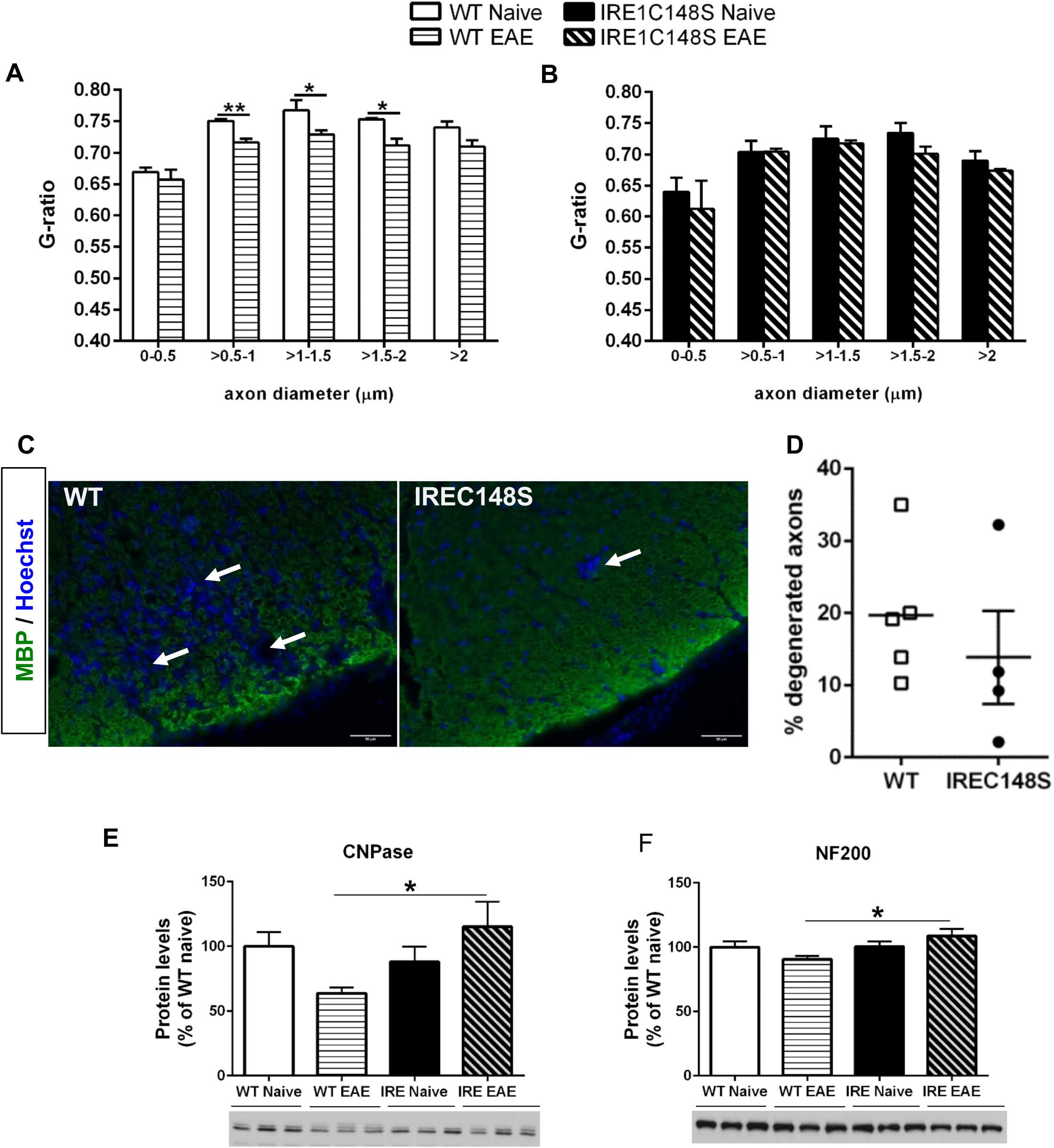
Reduction in myelin damage and reduced neuropathology in hyperactive IRE1C148S mice. **A**- G-ratio calculated in the lumbar spinal cord from WT mice naïve control (n=3) and 31d post-EAE (n=5). Bar graphs represent the mean ± sem. Multiple t-tests analysis, *p<0.05, **p<0.01. **B**- G-ratio calculated in the lumbar spinal cord from IRE1C148S mice naïve control (n=3) and 31d post-EAE (n=4). Bar graphs represent the mean ± sem. **C**- Micrographs of spinal cord from WT and IRE1C148S EAE mice (31d post immunization) immunostained for MBP. Hoechst is used as a nuclei counterstain. Scale bar: 50 μm. White arrows indicate areas of demyelination. **D**- Graph shows the percentage of degenerated axons in WT (n=5) and IRE1C148S (n=4) mice 31d post-EAE. **E-F**- Western blot analysis of lumbar spinal cord from WT and IRE1C148S naïve and 31 days post-EAE for myelin protein CNPase (**E**) and neuronal marker NF200 (**F**). Signal for each protein was normalized to the total protein amount quantified by Ponceau red staining of the membrane. Bar graphs represent the mean±sem of n=3/group. One Way ANOVA and Fisher’s test, *p<0.05.

### Increased MBP production but no difference in proliferation or differentiation of IRE1C148S mutant oligodendrocyte precursor cells

Since EAE is a demyelinating disease, and we found increased CNPase expression in IRE1C148S EAE compared to WT mice (Fig. 3D), we asked whether oligodendrocytes from IRE1C148S mutant mice were more efficient at making myelin proteins, *in vitro*. First, we compared the proliferative capabilities of both WT and IRE1C148S oligodendrocyte precursor cells (OPC) using EdU incorporation assay, and found no significant difference in the percentage of EdU^+^ cells, as shown in Fig. 4A-B. Next, we asked whether mutant IRE1 affected OPC maturation towards differentiaed oligodendrocytes. We cultured OPCs from WT and IRE1C148S, differentiated them for 5 days with thyroid hormone (T3), then evaluatedthe number of mature oligodendrocytes double labeled for CC1 and MBP. Neither the percentage of mature CC1^+^ oligodendrocytes (Fig. 4C) or CC1^+^MBP^+^ myelinating oligodendrocytes (Fig. 4D) exhibited a significant difference in IRE1C148S compared to WT cells, suggesting that the more active IRE1 does not affect the proliferation or differentiation of oligodendrocytes. Finally, we quantified the production of the myelin sheath, by integrating the intensity of MBP staining over each sheath area, and found an increase in myelin production in the IRE1C148S mutant mice (Fig. 4E). These results suggest that mutant oligodendrocytes produce more MBP than WT cells. Using two different inhibitors of IRE1α, A106 and 4μ8C, we confirmed that while OPCs still differentiate into mature oligodendrocytes (CC1+-cells), the production of myelin in culture is inhibited (Fig. 4F), suggesting a direct role for IRE1α in MBP expression and myelin production.

**Fig. 4.**
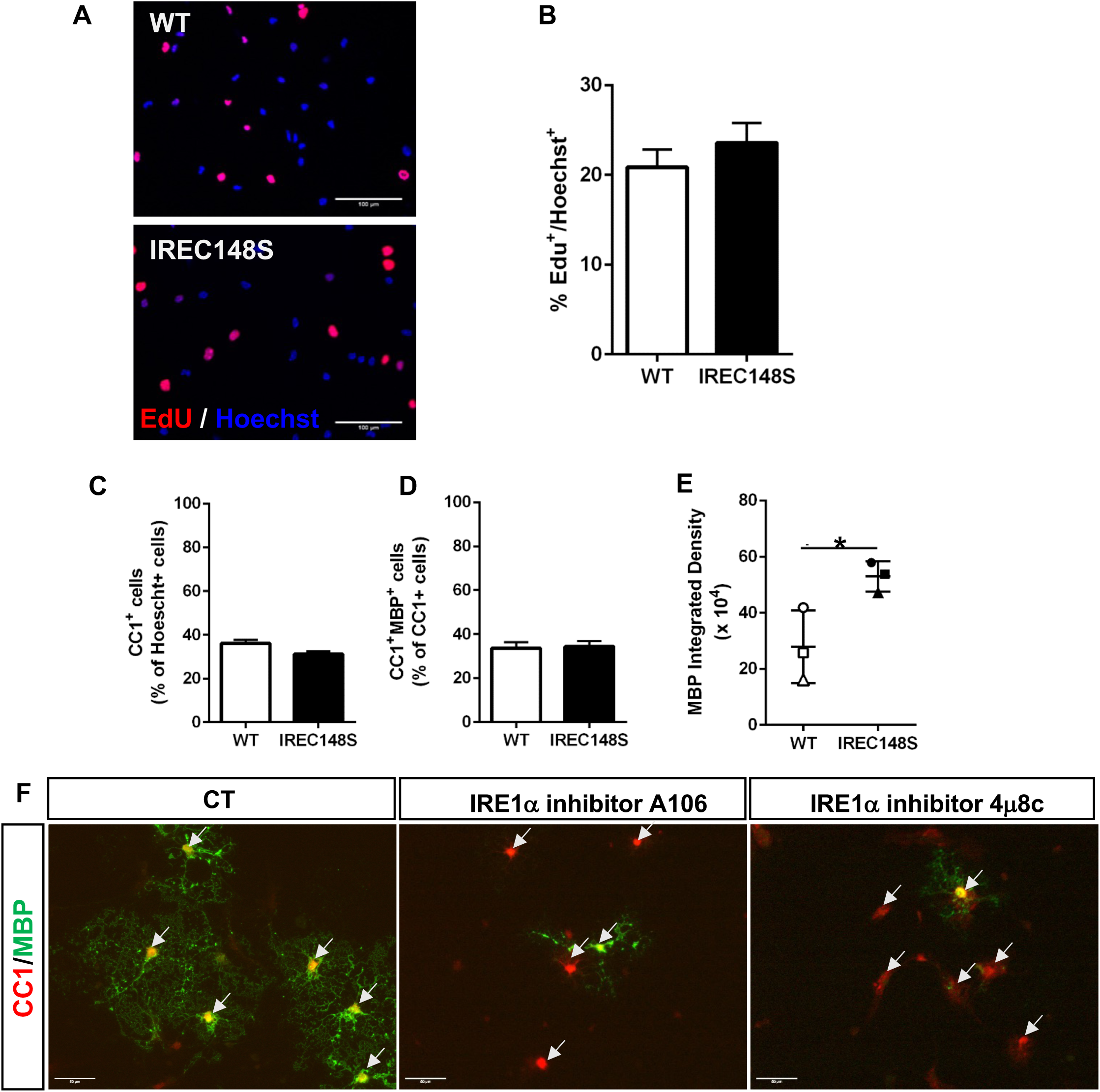
No difference in proliferation or differentiation rate of IRE1C148S mutant oligodendrocyte precursor cells but increase in MBP. **A**- Proliferation of oligodendrocytes precursor cell (OPC) from WT and IRE1C148S was determined by EdU incorporation. Representative images are shown. **B**- Quantification of the percentage of EdU^+^-cells in OPC cultures derived from WT and IRE1C148S. **C-D**-Quantification of the percentage of mature oligodendrocytes (CC1^+^-cells) between WT and IRE1C148S mutant (**C**), and the percentage of mature myelinating oligodendrocytes (CC1^+^MBP^+^, **D**). Bar graph represents the mean± sem of n=3/group. **E**- Quantification of the myelin sheath via MBP immunostaining. MBP integrated density was calculated for MBP^+^CC1^+^ oligodendrocytes from WT and IRE1C148S in triplicate. A paired T-test was used. *p<0.05 **F**- Effects of A106 and 4D8C, two inhibitors of IRE1α on OPC differentiation into myelinating (CC1^+^MBP^+^) oligodendrocytes. White arrows indicate CC1^+^ oligodendrocytes.

### Enhanced IRE1α activity reduces microgliosis and inflammatory gene expression

Neuroinflammation is a significant mediator of EAE disease pathology. Therefore, we investigated changes in spinal cord neuroinflammation in diseased WT and IRE1C148S mice. Astrocytes and microglia are two cell types that are activated under CNS pathologic conditions, with GFAP and IBA1 as their markers of reactivity, respectively. GFAP expression was upregulated in both WT and IRE1C148S EAE mice compared to their respective control group (Sup. Fig. 2). In contrast, reduced IBA1 upregulation was observed in diseased IRE1C148S mice (Fig. 5A), suggesting reduced activation of microglial cells in response to the disease. We next determined the levels of the proinflammatory molecules CCL2, TNF and IL1β in the spinal cords of 31d EAE mice. Gene expression of all three cytokines was reduced in diseased IRE1C148 mice in comparison to diseased WT mice (Fig. 5C-E). We also determined the levels of triggering receptor expressed on myeloid cells (TREM) 2, a receptor involved in phagocytosis, which were elevated in both diseased WT and IRE1C148S mice (Fig. 5F). Based on these data, we speculate that sustained IRE1 activity promotes reparative microglia resulting in attenuated neuroinflammation and enhanced phagocytosis.

**Fig. 5.**
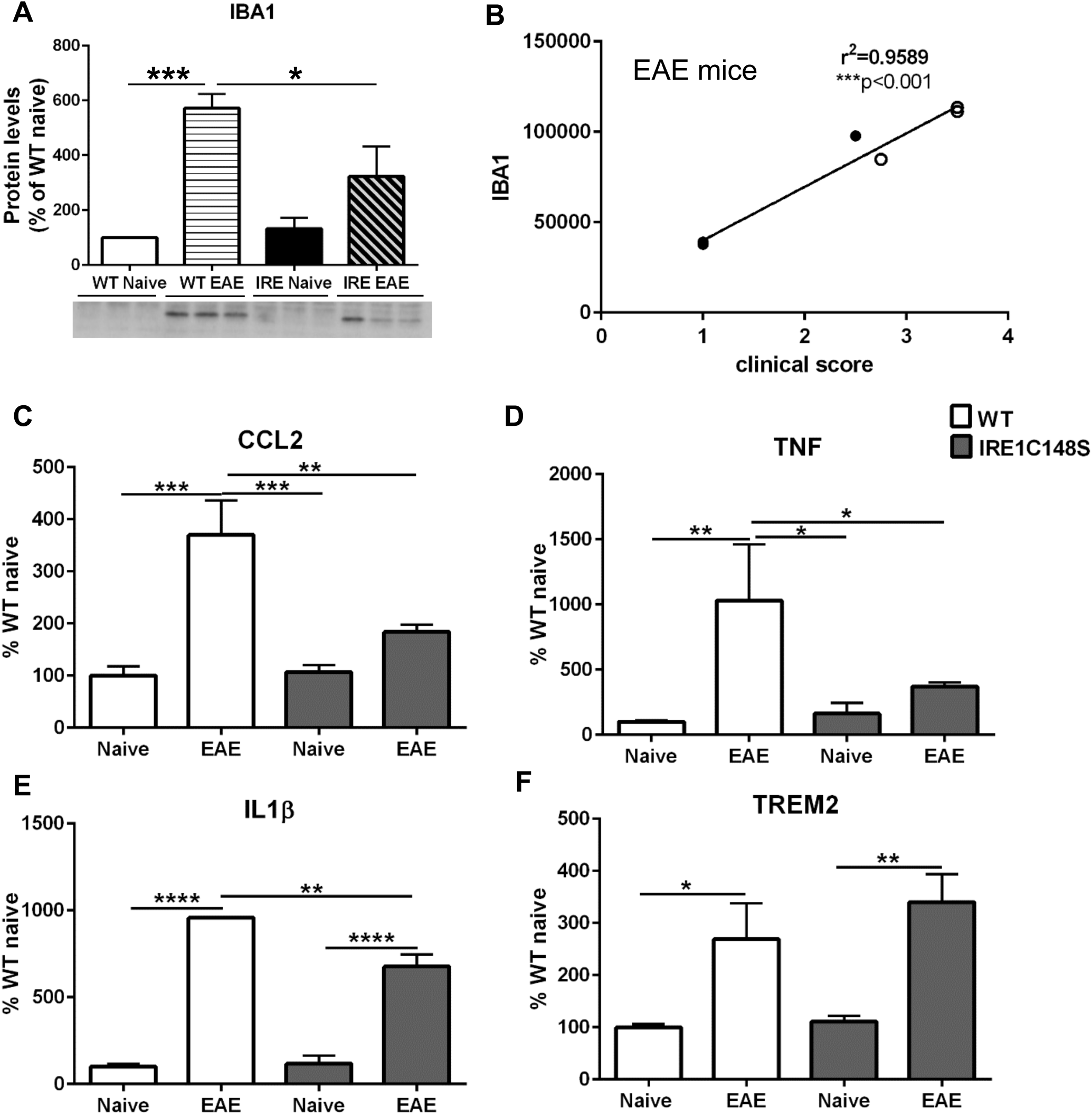
Reduction in proinflammatory cytokine levels but preservation of phagocytosis gene expression. **A-** Western blot analysis of lumbar spinal cord from WT and IRE1C148S naïve and 31 days post-EAE for IBA1, a marker of reactive microglia. Bar graph represents the mean±sem of n=3/group. *p<0.05, ***p<0.001 **B**- Correlation analysis between IBA1 protein levels in the spinal cord and the clinical score at d31 post-EAE. **C-F-** mRNA levels for proinflammatory cytokines CCL2 (**C**), TNF (**D**), IL1β (**E**), and TREM2, a marker of microglial phagocytosis (**F**), were determined in the lumbar spinal cord of WT and IRE1C148S naïve and 31d post EAE mice by quantitative real time PCR. Bar graphs represent the mean±SEM of n=3-5/group. *p<0.05, **p<0.01, “””p<0.001, ****p<0.0001

### Enhanced IRE1 activity in plasma cells does not increase antibody production

Apart from myelination, another example of dedicated high-volume secretion is production of antibodies by activation B cells. Therefore, we examined the antibody responses of IRE1C148S mice *in vitro* and *in vivo*. First, the mutation did not affect the development of a normal immune system, resulting in the expected subsets of splenic leukocytes (Sup. Fig. 3). Second, we asked whether differentiation of plasma cells, or their ability to secrete antibodies, were affected by the IRE1C148S mutation. Mice were immunized with NP-LPS and the appearance of plasma cells (B220^+^CD138^+^) in spleens 3 days post-injection was assessed by flow cytometric analysis. The absolute numbers of live plasma cells per organ and the percentages of plasma cells were slightly reduced in the IRE1C148S compared to WT (Fig. 6A). Purified splenic B cells were then stimulated *in vitro* with LPS plus IL5 for up to 72h. This treatment resulted in differentiation of B cells into plasma cells, as monitored by the percentage of CD138^+^ cells, but with no difference in the time course of differentiation whether the B cells carried WT or C148S IRE1 (Fig. 6) (Gaudette, Jones et al., 2020). Anti-NP antibody production by plasma cells was compared by ELISPOT assays, and both the frequency of antigen-specific cells and the appearance of their plaques were similar for both genotypes (Fig. 6C). Therefore, contrary to our expectation, hyperactive IRE1α does not enhance the differentiation of antibody producing cells nor does it increase antibody secretion *in vitro* or *in vivo*.

**Fig. 6.**
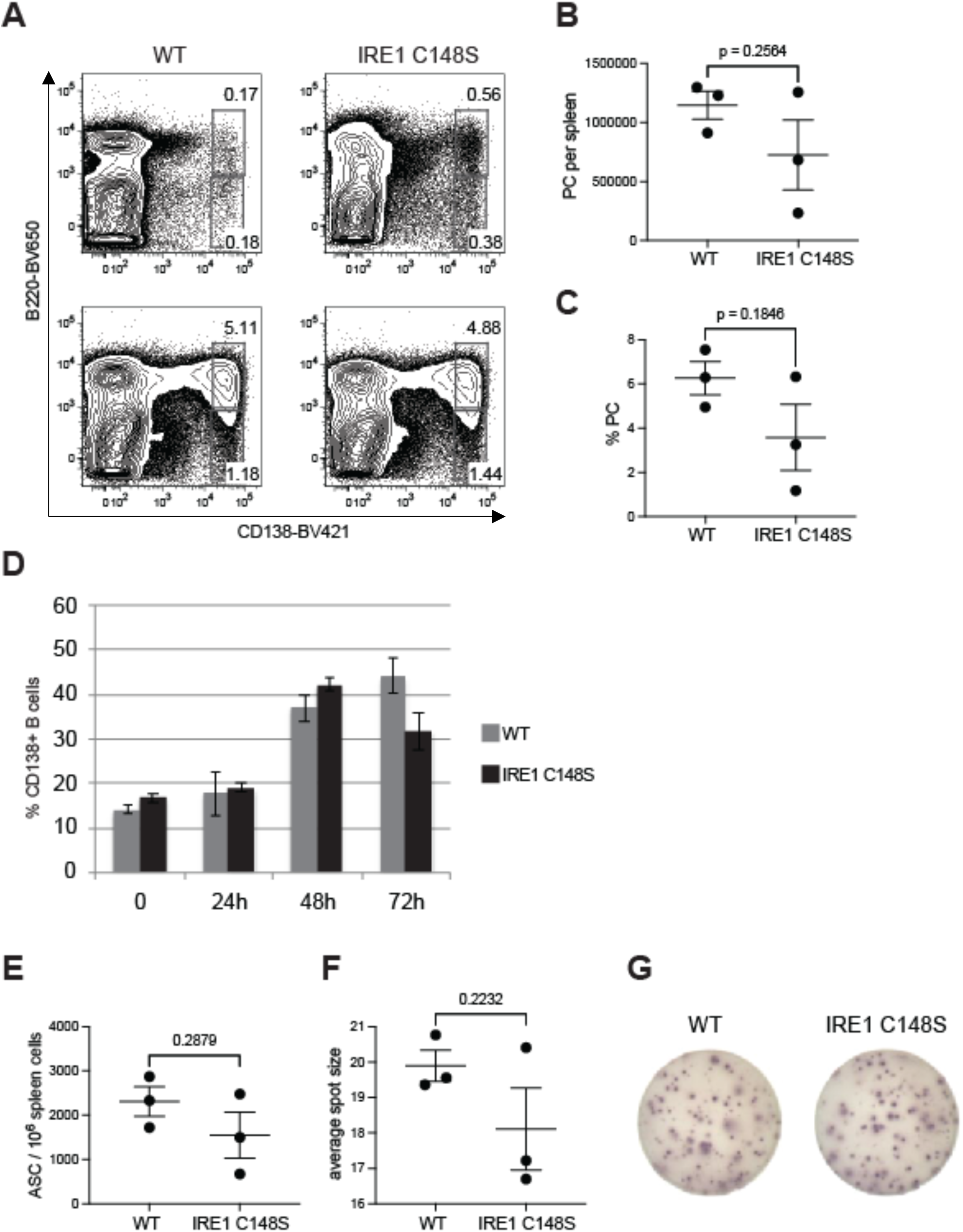
Hyperactive IRE1α does not increase production of antibodies by B-cell. **A-** Flow cytometric analysis of the differentiation of B-cells into plasma cells. Representative contour plots of splenocytes from WT and IRE1C148S mice, 72h after I.P. injection of NP-LPS (or Saline as a control). Plasma cells were gated as Dump [CD4,CD8a,Ter-119, F4/80]- IgD-, B220+/-, CD138+ cells. The top plots show steady-state controls and the bottom plots are the immunized mice. **B-C-** Quantification of antigen specific plasma cells (PC) in the spleens of WT and IRE1C148S mice (n=3 mice per group) B- Histogram of total number of NP-LPS-induced plasma cells (PC) in each spleen, C- The percentage of PC in each NP-LPS immunized spleen. **D-** Time course of *in vitro* differentiation of WT and IRE1C148S B cells into plasma cells following LPS and IL5 stimulation over 72h in culture. **E-** Anti-NP-LPS antibodies production assessed using ELISpot. The number of antibody secreting cells (ASC) from each spleen is shown as spots per million cells (n=3 mice per group) **F-** The amount of antibodies secreted from PC of each genotype is compared based of the calculated average spot size. **G-** Representative ELISpots for WT and IRE1C148S.

## Discussion

There is a considerable body of literature on the effects of decreasing or ablating IRE1α expression on the central and peripheral nervous system (23). On the other hand, there is very little information about the effects of upregulating IRE1α expression or activity, and this work describes a genetic approach to filling this information gap. We took advantage of our previous work in cell lines and primary cells in culture (14), and characterized a transgenic mouse model where an overactive IRE1α gene replaces the normal gene. The data presented in this manuscript shows that in two secretory cell types – myelin producing and antibody producing cells, increased IRE1α activity has different outcomes.

Our data show a remarkable protective effect of enhancing the activity of IRE1α on the progression of EAE, a mouse model of MS. Both the severity of motor defects and their duration are reduced. The most remarkable improvement is at the end of the time course (31d post-immunization), when disease progression seems to reverse consistently in IRE1C148S animals. Previous reports have demonstrated that IRE1α signaling plays an important role in development of the immune system and in immunological response to infection and disease (24). In particular, targeting the IRE1α gene (25) and its direct substrate XBP1 (26) have demonstrated that this signaling pathway is crucial for the development and function of antibody producing cells (27). Moreover, in addition to the XBP1 splicing activity of IRE1α, its RIDD activity is also involved in the differentiation of antibody secreting cells (28), suggesting that the quality of IRE1α output changes at various stages of differentiation (29). Because the IRE1α response is not constant, we asked if increased IRE1 activation further enhances either antibody production or plasma cell cellularity, using a genetic approach (14, 28). Surprisingly, our results demonstrate that there is no significant enhancement, indicating that the output from activation of wild type IRE1α in the B cell compartment is already optimal.

The dynamics of IRE1α activation have also been determined in models of inflammatory arthritis and EAE, though much less extensively. It was determined that deleting IRE1α in myeloid cells ameliorated chronic inflammation and disease pathology (30). Conversely, ectopic overexpression of spliced XBP1 with a viral vector was beneficial in a model of EAE/optic neuritis, when coupled with inhibition of PERK pathway (13), suggesting that activation of the IRE1α branch of UPR is protective but insufficient in this disease model. Therefore we used IRE1C148S mutation to determine how increased amplitude and duration of IRE1α activity affect EAE. In contrast to the effects on antibody secretion, we found that prolonging IRE1α activation improves functional outcome and reduces neuroinflammation and myelin damage in the EAE model. These results suggest a new interpretation to reports that luteolin or other flavonoids are therapeutic in models of EAE, spinal and brain injury (31–34): the positive therapeutic effects may be due to the ability of flavonoids to augment IRE1α activity (35) which then impact many other proteins. Indeed, luteolin is an IRE1 ligand in cells and in cell-free systems and it serves as an activator which bypasses the usual mode of stress-activated oligomerization (35–37). The increased splicing of XBP1 in IRE1C148S and in luteolin-treated cells may lead to transcriptional upregulation of target genes that are only moderately upregulated with the normal level of XBP1s in wild type cells.

IRE1α is unusual in that it possesses several enzymatic activities, each with outputs to different downstream pathways and distinct consequences. While we have only tested the XBP1 splicing activity of IRE1α, it is quite likely that other activities of IRE1α - higher RIDD activity and/or recruitment of JNK - are also affected by the C148S mutation. Increased spectrum of RIDD substrates is a particularly attractive and testable mode of output, since several transcripts with pro-apoptotic roles have already been shown to be cleaved by IRE1α (38–40). The marked differences in the effects on B lineage cells and CNS tissue could thus be attributed to the complexity of IRE1α signaling and to its differential regulation in various cell types, including immune cells (29, 41). Both deletion and robust pharmacological activation of IRE1α evidently impacts several critical signaling pathways including targets of XBP1s, of RIDD (42), JNK activation (11) and the mTOR pathway (28, 43) (28, 43), but the activation of these signaling pathways is likely different in distinct cell types.

The preservation of CNPase levels, along with decreased axonal degeneration and improved G-ratios in EAE animals carrying IRE1C148S mutation, indicate improved myelin integrity. Interestingly, as suggested by the findings of Unlu et al., CNPase itself could modulate the outcome of IRE1 activation (44). They demonstrated that CNPase is a component of the machinery that completes splicing of XBP1. Together with the RtcA gene product, CNPase acts to resolve the RNA end with 2’3’ cyclic phosphate, so that is can be ligated to 5’-OH RNA end by the ligase RtcB. In this manner, CNPase regulates the availability of spliced XBP1. Furthermore, the fine-tuning of XBP1 splicing by CNP and RtcA significantly alters the expression levels of direct targets of XBP1 (44). This may explain the involvement of CNPase in disease progression, as described in this work.

Microglial cells, also known as the resident macrophages of the brain, constantly ^i^monitor the CNS environment and respond to its perturbations. When they become inflammatory, they change their gene expression profile, including the upregulation of IBA1 (45) and of pro-inflammatory cytokines. We observed that a significant increase in IBA1 and in levels of several cytokines in WT EAE mice was markedly reversed in IREC148S EAE mice.

While IL1β gene expression was significantly reduced in IREC148S EAE mice compared to WT, the levels were still significantly higher than IREC148S naïve control mice and may be the reason why although there is improvement in motor function, mice still experience pain. It is also possible that increased IRE1α activity in peripheral leukocytes could lead to higher levels of prostaglandins and related pain, through XBP1s regulation of the prostaglandins synthase *Ptgs2* (46). The data also suggest that the pain and locomotor aspects of the disease are independent. Interestingly, in a EAE model where microglia are made more inflammatory at peak EAE by ablation of TNFR2, there is downregulation of many UPR genes, including XBP1 (supplementary Fig. 4, analysis of GEO set: GSE78082 from (47)), along with an upregulation of proinflammatory cytokines and a worse outcome for the mice. Our data show that the EAE IRE1C148S mutant mice with increased IRE1α activity and XBP1s levels have lower levels of inflammation, showing that IRE1C148S microglia are less inflammatory than WT microglia while still retaining their phagocytic activity.

Our mouse model has altered IRE1α activity in all cells and tissues, raising the question of which cell(s) is mostly accountable for the improved phenotype. Short of the definitive experiment of altering IRE1α in cell-specific fashion, we currently must rely on the indirect evidence to suggest that the most relevant cell type for IRE1 impact on EAE is microglia. While we see changes in neurons, oligodendrocytes and microglia, the decrease in pro-inflammatory cytokines and IBA1 levels in IREC148S EAE mice suggests a direct, cell-autonomous effect in the latter, whereas the improvements in neurons and oligodendrocytes may be either direct or secondary to decreased inflammation. Furthermore, the difference between cells that are affected by the hyperactive IRE1 and cells that are not, like plasma cells, is unexpected and fascinating. Microglia are not “professional” secretory cells like oligodendrocytes or plasma cells, which are programmed for high-rate production of myelin or immunoglobulin, respectively. Perhaps, they are not programmed to increase the secretory capacity by orders of magnitude, and therefore the impact of the overactive IRE1α is more pronounced in them.

At the molecular level, a likely explanation for cell-type different effects maybe cell specific interactome of IRE1α and unique XBP1s-regulated gene set expression (48). This is becoming more and more plausible, as there is an expanding number of IRE1α interactors that are being defined, such as the cytoskeleton-associated filamin A (49), TXNIP (50), and the TRAF6-mediated ubiquitination machinery (51).

The involvement of IRE1α in the inflammatory process in MS is not entirely unexpected. Prior studies in macrophages already showed that pro-inflammatory cytokine production is regulated (11, 52) by IRE1α activation as a consequence of TLR activation (30, 53). In that system, the activation of IRE1α is accompanied by activation of the TRAF2-ASK axis (54).

Taken together, the data suggest that activation of IRE1α signaling above and beyond the usual level is beneficial in this mouse model of MS, and that specific activators of IRE1α activity maybe useful tools in the treatment of MS.

## Supplementary Figure legends

**Supplementary Fig. 1.** Analysis of XBP1s expression levels in different brain regions (hippocampus, cortex and cerebellum).

**Supplementary Fig. 2.** Western blot analysis of the lumbar spinal cord of WT and IRE1C148S 31d post-EAE induction for GFAP. Data was normalized to GAPDH expression levels. Bar graphs represent the mean ± s.e.m. of n=3 per group.

**Supplementary Fig. 3.** Analysis of the different immune cell subsets in the spleens of WT and IRE1C148S mutant mice by flow cytometry shows no significant difference. Bars represent the mean ± s.e.m. of n=5 per group.

**Supplementary Fig. 4.** Analysis of GEO set: <u>GSE78082</u> (TNFR2-ablated microglia shows a more pro-inflammatory phenotype, with dysregulation of genes controlling innate immunity and host defense, (47)) for differentially expressed UPR genes. RNAseq was performed on WT and TNFR2^-/-^ microglia isolated from EAE mice.

## Supplementary methods

Flow cytometric analysis of the different cell subsets in the spleen. Adult WT and IREC148S mice (about 4-month-old) were euthanized by CO_2_ inhalation and spleens were quickly removed and placed in 2 ml of RPMI1640 on ice with ACK lysis buffer (Biolegend) to remove red blood cells. 2×10^6^ cells were stained using the Live/Dead™ aqua stain kit (Thermofisher Scientific) or Zombie Aqua live/dead kit (Biolegend) before incubation with the following antibody mixes (all antibodies were from Biolegend unless otherwise noted). Mix1 (B, T, NK cells): Alexa 488 anti-CD45 (clone), APC-Cy7 anti-CD19 (clone 6D5), BV650 anti-CD3 (clone 17A2), PE anti-CD4 (clone RM4-4), APC anti CD8 (clone 53-6.7) PE-Cy7 anti-NK1.1 (clone PK136). Mix2 (T-reg) Alexa 488 anti-CD45 (clone), BV650 anti-CD4 (clone RM4-5), APC anti-CD25 (clone PC61) and PE anti-Foxp3 (clone 150D). Mix3 (macrophages/monocytes, neutrophils) Alexa 488 anti-CD45 (clone), PE-Cy7 anti-CD11b (clone M1/70), PE anti-Ly6G (clone 1A8), APC-Cy7 anti-Ly6C (clone HK1.4). Plasma Cell panel: IgM-FITC Fab (Jackson Immunoresearch 115-097-020), CD138-BV421 (clone 281-2), B220-BV650 (clone RA3-6B2), CD19-BV785 (clone 6D5), CD38-AF700 (clone 90 eBioscience), IgD-APC-C7 (clone 11-26C.2A), CD4-PE-Cy (clone GK1.5), CD8a-PE-Cy7 (clone 53-6.7), Ter-119-PE-Cy7 (clone TER-119), F4/80-PE-Cy7 (clone BM8). In Mix2 cells were first stained with the surface antigens prior to fixation and permeabilization for Foxp3 intracellular staining using the True Nuclear™ transcription factor buffer set (Biolegend). Data were collected on a BD FACSymphony™ at the Flow Cytometry Facility of The Wistar Institute and analyzed with FlowJo v10.8 software (BD Life Sciences).

## Notes

### Competing Interest Statement

The authors have declared no competing interest.

## References

1. Cox, J. S., and Walter, P. (1996) A novel mechanism for regulating activity of a transcription factor that controls the unfolded protein response. Cell 87, 391–404

2. Uemura, A., Oku, M., Mori, K., and Yoshida, H. (2009) Unconventional splicing of XBP1 mRNA occurs in the cytoplasm during the mammalian unfolded protein response. J Cell Sci 122, 2877–2886

3. Lu, Y., Liang, F. X., and Wang, X. (2014) A synthetic biology approach identifies the mammalian UPR RNA ligase RtcB. Mol Cell 55, 758–770

4. Cox, J. S., Chapman, R. E., and Walter, P. (1997) The unfolded protein response coordinates the production of endoplasmic reticulum protein and endoplasmic reticulum membrane. Mol Biol Cell 8, 1805–1814

5. Stone, S., and Lin, W. (2015) The unfolded protein response in multiple sclerosis. Front Neurosci 9, 264

6. Scheper, W., and Hoozemans, J. J. (2015) The unfolded protein response in neurodegenerative diseases: a neuropathological perspective. Acta Neuropathol 130, 315–331

7. Fernandez, D., Geisse, A., Bernales, J. I., Lira, A., and Osorio, F. (2021) The Unfolded Protein Response in Immune Cells as an Emerging Regulator of Neuroinflammation. Frontiers in aging neuroscience 13, 682633

8. McGinley, A. M., Edwards, S. C., Raverdeau, M., and Mills, K. H. G. (2018) Th17cells, gammadelta T cells and their interplay in EAE and multiple sclerosis. J Autoimmun

9. Mhaille, A. N., McQuaid, S., Windebank, A., Cunnea, P., McMahon, J., Samali, A., and FitzGerald, U. (2008) Increased expression of endoplasmic reticulum stress-related signaling pathway molecules in multiple sclerosis lesions. J Neuropathol Exp Neurol 67, 200–211

10. Lin, W., and Popko, B. (2009) Endoplasmic reticulum stress in disorders of myelinating cells. Nat Neurosci 12, 379–385

11. Urano, F., Wang, X., Bertolotti, A., Zhang, Y., Chung, P., Harding, H. P., and Ron, D. (2000) Coupling of stress in the ER to activation of JNK protein kinases by transmembrane protein kinase IRE1. Science 287, 664–666

12. Lin, J. H., Li, H., Yasumura, D., Cohen, H. R., Zhang, C., Panning, B., Shokat, K. M., Lavail, M. M., and Walter, P. (2007) IRE1 signaling affects cell fate during the unfolded protein response. Science 318, 944–949

13. Huang, H., Miao, L., Liang, F., Liu, X., Xu, L., Teng, X., Wang, Q., Ridder, W. H., 3rd, Shindler, K. S., Sun, Y., and Hu, Y. (2017) Neuroprotection by eIF2alpha-CHOP inhibition and XBP-1 activation in EAE/optic neuritiss. Cell Death Dis 8, e2936

14. Eletto, D., Eletto, D., Dersh, D., Gidalevitz, T., and Argon, Y. (2014) Protein disulfide isomerase A6 controls the decay of IRE1alpha signaling via disulfide-dependent association. Molecular cell 53, 562–576

15. Murphy, K. L., Fischer, R., Swanson, K. A., Bhatt, I. J., Oakley, L., Smeyne, R., Bracchi-Ricard, V., and Bethea, J. R. (2020) Synaptic alterations and immune response are sexually dimorphic in a non-pertussis toxin model of experimental autoimmune encephalomyelitis. Exp Neurol 323, 113061

16. Krementsov, D. N., Noubade, R., Dragon, J. A., Otsu, K., Rincon, M., and Teuscher, C. (2014) Sex-specific control of central nervous system autoimmunity by p38 mitogen-activated protein kinase signaling in myeloid cells. Ann Neurol 75, 50–66

17. Fischer, R., Padutsch, T., Bracchi-Ricard, V., Murphy, K. L., Martinez, G. F., Delguercio, N., Elmer, N., Sendetski, M., Diem, R., Eisel, U. L. M., Smeyne, R. J., Kontermann, R. E., Pfizenmaier, K., and Bethea, J. R. (2019) Exogenous activation of tumor necrosis factor receptor 2 promotes recovery from sensory and motor disease in a model of multiple sclerosis. Brain Behav Immun 81, 247–259

18. Brambilla, R., Morton, P. D., Ashbaugh, J. J., Karmally, S., Lambertsen, K. L., and Bethea, J. R. (2014) Astrocytes play a key role in EAE pathophysiology by orchestrating in the CNS the inflammatory response of resident and peripheral immune cells and by suppressing remyelination. Glia 62, 452–467

19. Gaudette, B. T., Jones, D. D., Bortnick, A., Argon, Y., and Allman, D. (2020) mTORC1 coordinates an immediate unfolded protein response-related transcriptome in activated B cells preceding antibody secretion. Nature communications 11, 723

20. Jones, D. D., Gaudette, B. T., Wilmore, J. R., Chernova, I., Bortnick, A., Weiss, B. M., and Allman, D. (2016) mTOR has distinct functions in generating versus sustaining humoral immunity. The Journal of clinical investigation 126, 4250–4261

21. Chapman, R., Sidrauski, C., and Walter, P. (1998) Intracellular signaling from the endoplasmic reticulum to the nucleus. Annu Rev Cell Dev Biol 14, 459–485

22. Madsen, P. M., Motti, D., Karmally, S., Szymkowski, D. E., Lambertsen, K. L., Bethea, J. R., and Brambilla, R. (2016) Oligodendroglial TNFR2 Mediates Membrane TNF-Dependent Repair in Experimental Autoimmune Encephalomyelitis by Promoting Oligodendrocyte Differentiation and Remyelination. The Journal of neuroscience : the official journal of the Society for Neuroscience 36, 5128–5143

23. Ni, H., Rui, Q., Li, D., Gao, R., and Chen, G. (2018) The Role of IRE1 Signaling in the Central Nervous System Diseases. Curr Neuropharmacol 16, 1340–1347

24. So, J. S. (2018) Roles of Endoplasmic Reticulum Stress in Immune Responses. Mol Cells 41, 705–716

25. Tufanli, O., Telkoparan Akillilar, P., Acosta-Alvear, D., Kocaturk, B., Onat, U. I., Hamid, S. M., Cimen, I., Walter, P., Weber, C., and Erbay, E. (2017) Targeting IRE1 with small molecules counteracts progression of atherosclerosis. Proceedings of the National Academy of Sciences of the United States of America 114, E1395–E1404

26. Hu, C. C., Dougan, S. K., McGehee, A. M., Love, J. C., and Ploegh, H. L. (2009) XBP-1 regulates signal transduction, transcription factors and bone marrow colonization in B cells. The EMBO journal 28, 1624–1636

27. Iwakoshi, N. N., Lee, A. H., Vallabhajosyula, P., Otipoby, K. L., Rajewsky, K., and Glimcher, L. H. (2003) Plasma cell differentiation and the unfolded protein response intersect at the transcription factor XBP-1. Nat Immunol 4, 321–329

28. Benhamron, S., Hadar, R., Iwawaky, T., So, J. S., Lee, A. H., and Tirosh, B. (2014) Regulated IRE1-dependent decay participates in curtailing immunoglobulin secretion from plasma cells. European journal of immunology 44, 867–876

29. Ricci, D., Gidalevitz, T., and Argon, Y. (2021) The special unfolded protein response in plasma cells. Immunol Rev 303, 35–51

30. Qiu, Q., Zheng, Z., Chang, L., Zhao, Y. S., Tan, C., Dandekar, A., Zhang, Z., Lin, Z., Gui, M., Li, X., Zhang, T., Kong, Q., Li, H., Chen, S., Chen, A., Kaufman, R. J., Yang, W. L., Lin, H. K., Zhang, D., Perlman, H., Thorp, E., Zhang, K., and Fang, D. (2013) Toll-like receptor-mediated IRE1alpha activation as a therapeutic target for inflammatory arthritis. The EMBO journal 32, 2477–2490

31. Verbeek, R., van Tol, E. A., and van Noort, J. M. (2005) Oral flavonoids delay recovery from experimental autoimmune encephalomyelitis in SJL mice. Biochemical pharmacology 70, 220–228

32. Theoharides, T. C. (2009) Luteolin as a therapeutic option for multiple sclerosis. J Neuroinflammation 6, 29

33. Xu, J., Wang, H., Ding, K., Zhang, L., Wang, C., Li, T., Wei, W., and Lu, X. (2014) Luteolin provides neuroprotection in models of traumatic brain injury via the Nrf2-ARE pathway. Free radical biology & medicine 71, 186–195

34. Kempuraj, D., Thangavel, R., Kempuraj, D. D., Ahmed, M. E., Selvakumar, G. P., Raikwar, S. P., Zaheer, S. A., Iyer, S. S., Govindarajan, R., Chandrasekaran, P. N., and Zaheer, A. (2021) Neuroprotective effects of flavone luteolin in neuroinflammation and neurotrauma. BioFactors 47, 190–197

35. Wiseman, R. L., Zhang, Y., Lee, K. P., Harding, H. P., Haynes, C. M., Price, J., Sicheri, F., and Ron, D. (2010) Flavonol activation defines an unanticipated ligand-binding site in the kinase-RNase domain of IRE1. Mol Cell 38, 291–304

36. Meares, G. P., Hughes, K. J., Naatz, A., Papa, F. R., Urano, F., Hansen, P. A., Benveniste, E. N., and Corbett, J. A. (2011) IRE1-dependent activation of AMPK in response to nitric oxide. Molecular and cellular biology 31, 4286–4297

37. Ricci, D., Marrocco, I., Blumenthal, D., Dibos, M., Eletto, D., Vargas, J., Boyle, S., Iwamoto, Y., Chomistek, S., Paton, J. C., Paton, A. W., and Argon, Y. (2019) Clustering of IRE1alpha depends on sensing ER stress but not on its RNase activity. FASEB J 33, 9811–9827

38. Coelho, D. S., and Domingos, P. M. (2014) Physiological roles of regulated Ire1 dependent decay. Frontiers in genetics 5, 76

39. Groenendyk, J., Peng, Z., Dudek, E., Fan, X., Mizianty, M. J., Dufey, E., Urra, H., Sepulveda, D., Rojas-Rivera, D., Lim, Y., Kim, D. H., Baretta, K., Srikanth, S., Gwack, Y., Ahnn, J., Kaufman, R. J., Lee, S. K., Hetz, C., Kurgan, L., and Michalak, M. (2014) Interplay between the oxidoreductase PDIA6 and microRNA-322 controls the response to disrupted endoplasmic reticulum calcium homeostasis. Sci Signal 7, ra54

40. Hussien, Y., Cavener, D. R., and Popko, B. (2014) Genetic inactivation of PERK signaling in mouse oligodendrocytes: normal developmental myelination with increased susceptibility to inflammatory demyelination. Glia 62, 680–691

41. Osorio, F., Tavernier, S. J., Hoffmann, E., Saeys, Y., Martens, L., Vetters, J., Delrue, I., De Rycke, R., Parthoens, E., Pouliot, P., Iwawaki, T., Janssens, S., and Lambrecht, B. N. (2014) The unfolded-protein-response sensor IRE-1alpha regulates the function of CD8alpha+ dendritic cells. Nat Immunol 15, 248–257

42. Tang, C. H., Chang, S., Paton, A. W., Paton, J. C., Gabrilovich, D. I., Ploegh, H. L., Del Valle, J. R., and Hu, C. C. (2018) Phosphorylation of IRE1 at S729 regulates RIDD in B cells and antibody production after immunization. The Journal of cell biology 217, 1739–1755

43. Benhamron, S., Pattanayak, S. P., Berger, M., and Tirosh, B. (2015) mTOR activation promotes plasma cell differentiation and bypasses XBP-1 for immunoglobulin secretion. Molecular and cellular biology 35, 153–166

44. Unlu, I., Lu, Y., and Wang, X. (2018) The cyclic phosphodiesterase CNP and RNA cyclase RtcA fine-tune noncanonical XBP1 splicing during ER stress. The Journal of biological chemistry 293, 19365–19376

45. Imai, Y., and Kohsaka, S. (2002) Intracellular signaling in M-CSF-induced microglia activation: role of Iba1. Glia 40, 164–174

46. Chopra, S., Giovanelli, P., Alvarado-Vazquez, P. A., Alonso, S., Song, M., Sandoval, T. A., Chae, C. S., Tan, C., Fonseca, M. M., Gutierrez, S., Jimenez, L., Subbaramaiah, K., Iwawaki, T., Kingsley, P. J., Marnett, L. J., Kossenkov, A. V., Crespo, M. S., Dannenberg, A. J., Glimcher, L. H., Romero-Sandoval, E. A., and Cubillos-Ruiz, J. R. (2019) IRE1alpha-XBP1 signaling in leukocytes controls prostaglandin biosynthesis and pain. Science 365

47. Gao, H., Danzi, M. C., Choi, C. S., Taherian, M., Dalby-Hansen, C., Ellman, D. G., Madsen, P. M., Bixby, J. L., Lemmon, V. P., Lambertsen, K. L., and Brambilla, R. (2017) Opposing Functions of Microglial and Macrophagic TNFR2 in the Pathogenesis of Experimental Autoimmune Encephalomyelitis. Cell reports 18, 198–212

48. Park, S. M., Kang, T. I., and So, J. S. (2021) Roles of XBP1s in Transcriptional Regulation of Target Genes. Biomedicines 9

49. Urra, H., Henriquez, D. R., Canovas, J., Villarroel-Campos, D., Carreras-Sureda, A., Pulgar, E., Molina, E., Hazari, Y. M., Limia, C. M., Alvarez-Rojas, S., Figueroa, R., Vidal, R. L., Rodriguez, D. A., Rivera, C. A., Court, F. A., Couve, A., Qi, L., Chevet, E., Akai, R., Iwawaki, T., Concha, M. L., Glavic, A., Gonzalez-Billault, C., and Hetz, C. (2018) IRE1alpha governs cytoskeleton remodelling and cell migration through a direct interaction with filamin A. Nature cell biology 20, 942–953

50. Oslowski, C. M., Hara, T., O’Sullivan-Murphy, B., Kanekura, K., Lu, S., Hara, M., Ishigaki, S., Zhu, L. J., Hayashi, E., Hui, S. T., Greiner, D., Kaufman, R. J., Bortell, R., and Urano, F. (2012) Thioredoxin-interacting protein mediates ER stress-induced beta cell death through initiation of the inflammasome. Cell metabolism 16, 265–273

51. Deng, L., Wang, C., Spencer, E., Yang, L., Braun, A., You, J., Slaughter, C., Pickart, C., and Chen, Z. J. (2000) Activation of the IkappaB kinase complex by TRAF6 requires a dimeric ubiquitin-conjugating enzyme complex and a unique polyubiquitin chain. Cell 103, 351–361

52. Luo, D., He, Y., Zhang, H., Yu, L., Chen, H., Xu, Z., Tang, S., Urano, F., and Min, W. (2008) AIP1 is critical in transducing IRE1-mediated endoplasmic reticulum stress response. The Journal of biological chemistry 283, 11905–11912

53. Martinon, F., and Glimcher, L. H. (2011) Regulation of innate immunity by signaling pathways emerging from the endoplasmic reticulum. Current opinion in immunology 23, 35–40

54. Matsuzawa, A., and Ichijo, H. (2008) Redox control of cell fate by MAP kinase: physiological roles of ASK1-MAP kinase pathway in stress signaling. Biochim Biophys Acta 1780, 1325–1336

55. Volmer, R., van der Ploeg, K., and Ron, D. (2013) Membrane lipid saturation activates endoplasmic reticulum unfolded protein response transducers through their transmembrane domains. Proc Natl Acad Sci U S A 110, 4628–4633

