## Supplementary figures and images for "Increased activity of IRE1 improves the clinical presentation of EAE"

### Supplemental Figures

**Supplementary Figure 1**

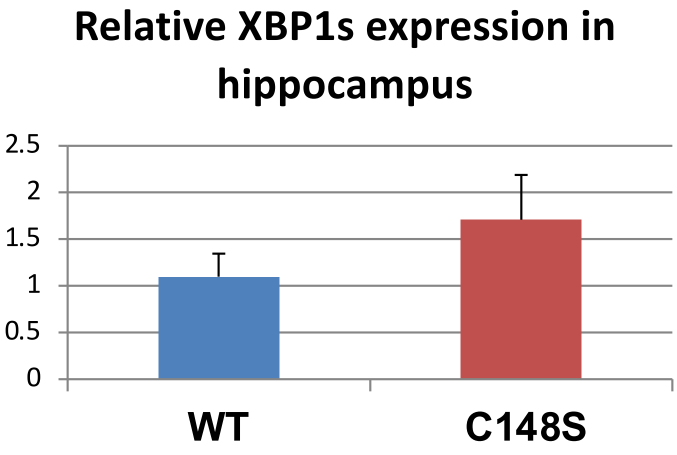


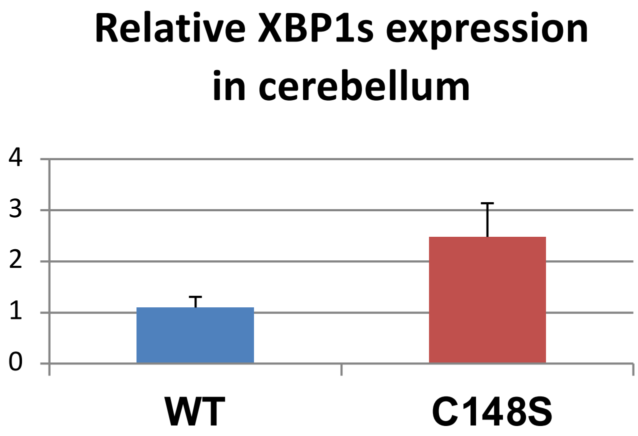


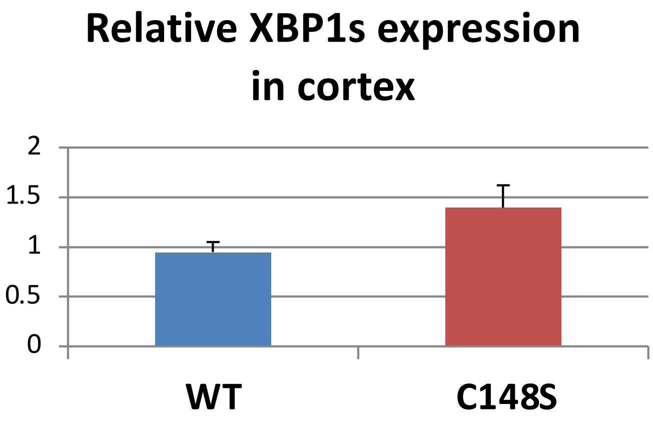

**Supplementary Figure 2**

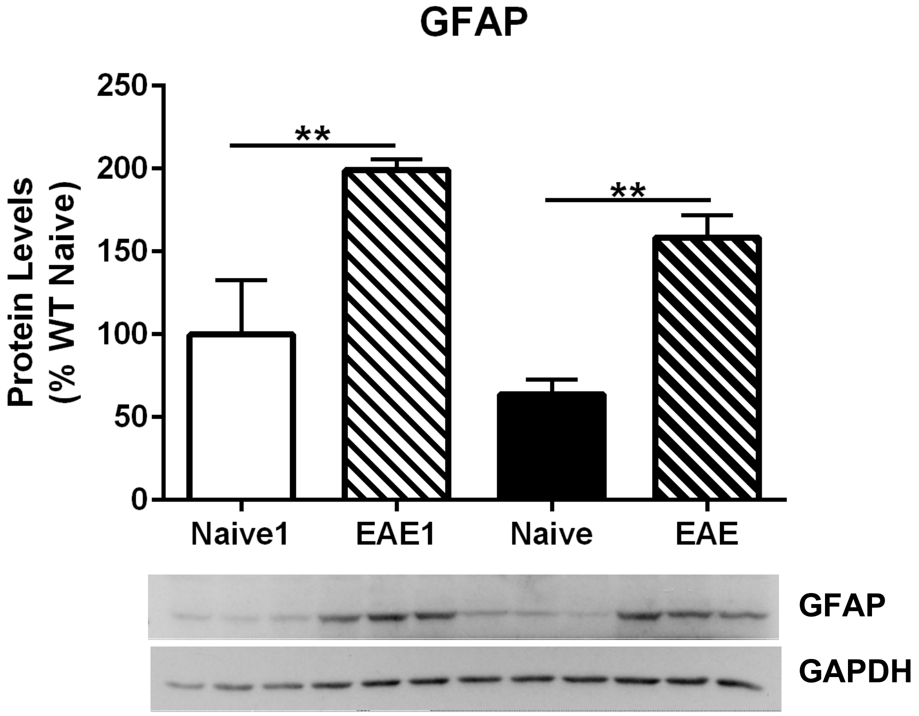

#### Supplementary Figure 3

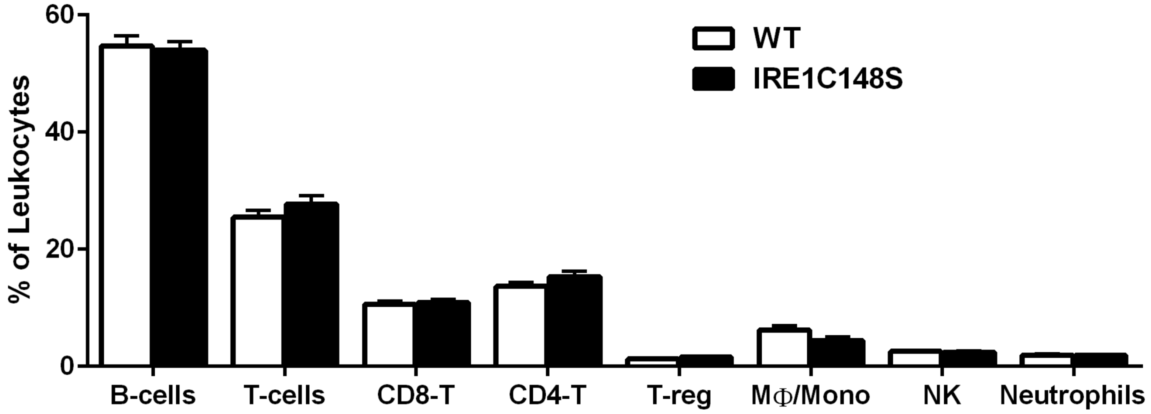

**Supplementary Figure 4**

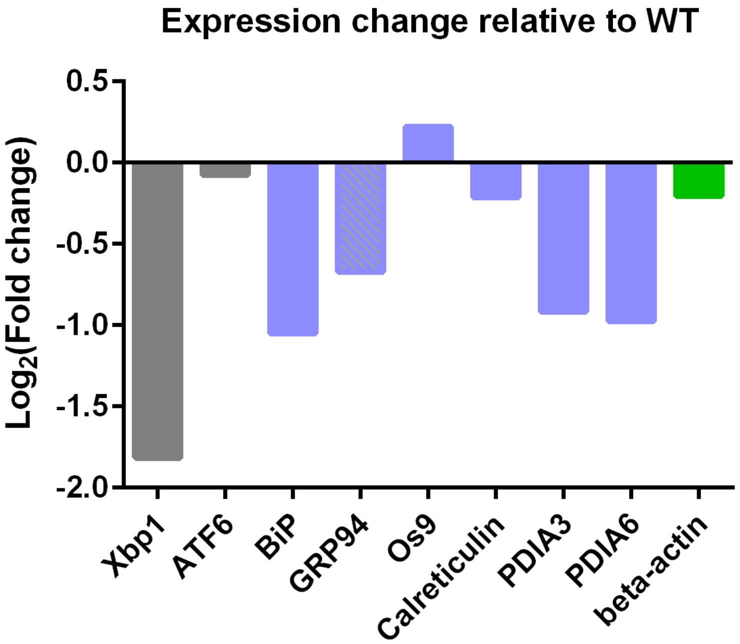
